# What’s in Your Gamma? Activation of The Ventral Fronto-Parietal Attentional Network in Response to Distracting Sounds

**DOI:** 10.1101/469783

**Authors:** Hesham A. ElShafei, Lesly Fornoni, Rémy Masson, Olivier Bertrand, Aurélie Bidet-Caulet

## Abstract

Auditory attention operates through top-down (TD) and bottom-up (BU) mechanisms that are supported by dorsal and ventral brain networks, respectively, with the main overlap in the lateral prefrontal cortex (lPFC). A good TD/BU balance is essential to be both task-efficient and aware of our environment, yet it is rarely investigated. Oscillatory activity is a novel method to probe the attentional dynamics with evidence that gamma activity (>30Hz) could signal BU processing and thus would be a good candidate to support the activation of the ventral BU network. MEG data were collected from 21 young adults performing the Competitive Attention Task which enables simultaneous investigation of BU and TD attentional mechanisms. Distracting sounds elicited an increase in gamma activity in regions of the BU ventral network. TD attention modulated these gamma responses in regions of the inhibitory cognitive control system: the medial prefrontal and anterior cingulate cortices. Finally, distracting-sound-induced gamma activity was synchronous between the auditory cortices and several distant brain regions, notably the lPFC. We provide novel insight into the role of gamma activity (1) in supporting the activation of the BU ventral network, and (2) in subtending the TD/BU attention balance in the prefrontal cortex.

## INTRODUCTION

In an environment that contains far more information than we can process at a time, we rely on our attention to prioritize the processing of only a fragment of these incoming stimuli (Desimone and Duncan 1995). Attention can be oriented endogenously (top-down), in anticipation of an upcoming stimulus for example, or it can be captured exogenously (bottom-up) by a salient irrelevant stimulus such as a telephone ringing (Posner and Petersen 1990; Petersen and Posner 2012). A dynamic balance between top-down (TD) and bottom-up (BU) mechanisms of attention is essential to be task-efficient while being aware, yet not fully distracted, of our surroundings.

Two major neural networks support TD (endogenous) and BU (exogenous) mechanisms of attention: a dorsal frontoparietal network comprising the posterior frontal and intraparietal cortices; and a ventral frontoparietal network, largely lateralized to the right hemisphere, comprising the temporo-parietal junction (TPJ) and the ventral prefrontal cortex (vPFC); with the two networks overlapping mainly in the lateral prefrontal cortex (lPFC) (Kim et al. 1999; Miller and Cohen 2001; Corbetta and Shulman 2002; Fox et al. 2006; He et al. 2007; Corbetta et al. 2008; Asplund et al. 2010).

A promising way of addressing the dynamics of TD and BU attentional systems is to explore brain rhythms. On one hand, TD anticipatory attention is indexed by (de)synchronisation of oscillatory activity in the alpha band (review in Klimesch et al. 2007; Jensen and Mazaheri 2010; Frey et al. 2014; e.g. ElShafei et al. 2018). On the other hand, BU attentional capture is signalled by an increased activity in the gamma band in the primate brain (Buschman and Miller, 2007).

Activation in the gamma band (>30 Hz) has been associated to attention, with enhanced gamma activity in the visual (e.g. Fries et al. 2001) or auditory (e.g. Ray et al. 2008) cortices, in response to attended visual or auditory stimuli, respectively. Gamma activity has also been observed in regions other than sensory regions (e.g. dorsolateral prefrontal cortex, intraparietal sulcus and temporo-parietal junction) in several working memory (Michels et al. 2010; Albouy et al. 2013), and visual (Akimoto et al. 2013, 2014) or auditory (Lee et al. 2007; Ahveninen et al. 2013) oddball tasks. Thus, gamma activity seems to promote the activation of relevant processes across the brain and not only in sensory cortices. Finally, it has been demonstrated that attention increases the coupling between frontal and relevant sensory regions via gamma synchrony (Buschman and Miller 2007; Gregoriou et al. 2009; Baldauf and Desimone 2014). Therefore, gamma activity would be a good candidate to support activation of the ventral network of attention. However, to our knowledge, no study has investigated the role of gamma activity in the balance between BU and TD mechanisms of attention in the human brain.

Distraction by unexpected sounds has been mostly investigated using variations of the reaction-time based oddball paradigm. In this paradigm, the response time to a target stimulus is compared in the presence (vs. absence) of a rare deviation (oddball) within a sequence of irrelevant stimuli that could be in the same (Schröger 1996) or a different (Escera et al. 1998) modality than the attended target. However, the adequacy of such paradigm to provide a reliable measure of attentional capture has been recently criticized (review in Parmentier and Andrés 2010; Bidet-Caulet et al. 2014; Dalton and Hughes 2014; Masson and Bidet-Caulet 2018).

In 2014, Bidet-Caulet and colleagues proposed a novel paradigm, the Competitive Attention Task (CAT), an adaptation of the Posner cueing paradigm using visual cues and monaural auditory targets. In this task, TD anticipatory attention is measured by comparing trials with informative cues to trials with uninformative cues. BU attentional capture is triggered by a binaural distracting sound played during the delay between the cue and the target in only 25 % of the trials. Distraction is assessed as the impact of distracting sounds on task performance and the balance between TD and BU mechanisms can be measured by comparing responses to distracting sounds following informative vs. uninformative cues.

We have recorded MEG activity from young healthy adults performing the CAT to test the following hypotheses. (1) BU attentional capture by an isolated unexpected stimulus would be indexed by gamma activity in the ventral attentional network including the TPJ and vPFC. (2) The lPFC would support the balance between BU and TD attention by demonstrating modulations of gamma activity to distracting sounds by cue information, since the lPFC is part of both the ventral and dorsal networks of attention. Finally, we sought to investigate the connectivity, subtended by gamma activity, between the auditory cortices and other brain regions during the presentation of a distracting sound, with a prediction that the mains hubs of this connectivity would lie within the lPFC.

## 1 MATERIAL & METHODS

### 1.1 Participants

Twenty-one healthy participants (9 females) took part in this study. The mean age was 24.7 years ± 0.62 Standard Error of Mean (SEM). All participants were right handed, and reported normal hearing, and normal or corrected-to-normal vision. All participants were free from any neurological or psychiatric disorders. The study was approved by the local ethical committee, and subjects gave written informed consent, according to the Declaration of Helsinki, and they were paid for their participation.

### 1.2 Stimuli and tasks

#### 1.2.1 Competitive Attention Task (CAT)

In 75 % of the trials, a target sound (100 ms duration) followed a central visual cue (200 ms duration) with a fixed delay of 1000 ms (see Figure 1). The cue was a green arrow, presented on a grey-background screen, pointing either to the left, right, or both sides. Target sounds were monaural pure tones (carrier frequency between 512 and 575 Hz; 5 ms rise-time, 5 ms fall-time). In the other 25 %, the same structure was retained, however, a binaural distracting sound (300 ms duration) was played during the cue-target delay (50-650 ms range after cue offset). Trials with a distracting sound played from 50 ms to 350 ms after the cue offset were classified as DIS1, those with a distracting sound played from 350 ms to 650 ms after the cue offset were classified as DIS2, those with no distracting sound were classified as NoDIS. A total of 40 different ringing sounds were used as distracting sounds (clock-alarm, door-bell, phone ring, etc.) for each participant.

**Figure 1.**
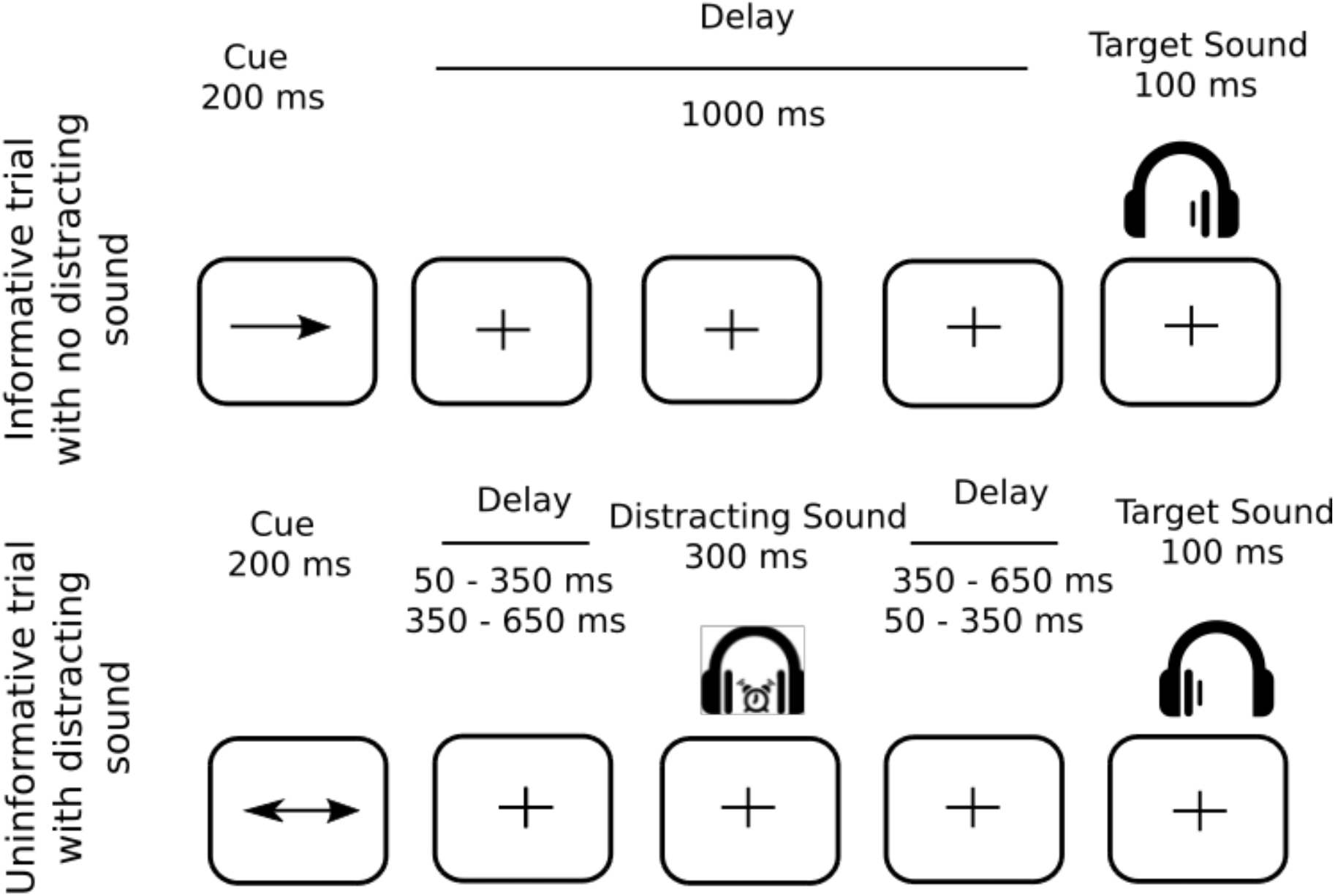
Protocol. Top row. Example of an informative trial with no distracting sound: a one-sided visual cue (200 ms duration) indicated in which ear (left or right) the target sound would be played (100 ms duration) after a fixed 1000-ms delay. Bottom row. Example of an uninformative trial with a distracting sound: a two-sided visual cue (200 ms duration) did not provide any indication in which ear (left or right) the target sound will be played. In 25 % of the trials, a binaural distracting sound (300 ms duration), such as a clock ring, was played during the delay between cue and target. The distracting sound could equiprobably onset in two different time periods after the cue offset: in the 50–350 ms range, or in the 350–650 ms range.

The cue and target categories were manipulated in the same proportion for trials with and without distracting sound. In 25% of the trials, the cue was pointing left, and the target sound was played in the left ear, and in 25% of the trials, the cue was pointing right, and the target sound was played in the right ear, leading to a total of 50% of *informative* trials. In the other 50% of the trials, the cue was *uninformative*, pointing in both directions, and the target sound was played in the left (25%) or right (25%) ear. To compare brain responses to acoustically matched sounds, the same distracting sounds were played in each combination of cue category (informative, uninformative) and distractor condition (DIS1 or DIS2). Each distracting sound was thus played 4 times during the whole experiment, but no more than once during each single block to limit habituation. Participants were instructed to categorize two target sounds as either high- or low-pitched sound, by either pulling or pushing a joystick.

The target type (high or low) was manipulated in the same proportion in all conditions. The mapping between the targets (low or high) and the responses (pull or push) was counterbalanced across participants but did not change across the blocks for each participant. In order to account for the participants’ pitch-discrimination capacities, the pitch difference between the two target sounds was defined in a Discrimination Task (see below). Participants were informed that informative cues were 100 % predictive and that a distracting sound could be sometimes played. They were asked to allocate their attention to the cued side in the case of informative cue, to ignore the distractors and to respond as quickly and correctly as possible. Participants had a 3.4 second (3400 ms) response window. In the absence of the visual cue, a blue fixation cross was presented at the center of the screen. Subjects were instructed to keep their eyes fixated on the cross.

#### 1.2.2 Discrimination Task

Participants were randomly presented with one of two target sounds: a low-pitched sound (512 Hz) and a high-pitched sound (575 Hz; two semitones higher), equiprobably in each ear (four trials per ear and per pitch). As described above, participants were asked to categorize the target sounds as either high- or low-pitched sound within 3 seconds.

#### 1.2.3 Procedure

Participants were seated in a sound-attenuated, magnetically shielded recording room, at a 50 cm distance from the screen. The response device was an index-operated joystick that participants moved either towards them (when instructed to pull) or away from them (when instructed to push). All stimuli were delivered using Presentation software (Neurobehavioral Systems, Albany, CA, USA). All sounds were presented through air-conducting tubes using Etymotic ER-3A foam earplugs (Etymotic Research, Inc., USA).

First, the auditory threshold was determined for the two target sounds differing by 2 semitones (512 and 575 Hz), for each ear, for each participant using the Bekesy tracking method (Von Békésy and Wever 1960). The target sounds were then monaurally presented at 25 dB sensation level (between 37.5 and 52.1 dB A across subjects) while the distracting sounds were binaurally played at 55 dB sensation level (between 47.5 and 62.1 dB A across subjects), above the target sound thresholds. Second, participants performed the Discrimination task. Afterwards, participants were trained with a short sequence of the Competitive Attention Task. Finally, MEG and EEG were recorded while subjects performed 10 blocks (64 trials each) leading to 240 trials in the NoDIS and 80 in the DIS conditions, for informative and uninformative cues, separately. The whole recording session lasted around 80 minutes. After the MEG/EEG session, participants’ subjective reports regarding their strategies were collected.

### 1.3 Behavioral Data Analysis

For behavioral data analysis, a button press before target onset was considered as a false alarm (FA). A trial with no button-press after target onset and before the next cue onset was considered as a miss trial. A trial with no FA and with a button-press after target onset was counted as correct if the pressed button matched the response mapped to the target sound, and as incorrect if otherwise. Reaction-times (RTs) to targets were analysed in the correct trials only.

The influence of (1) cue condition (2 levels: informative and uninformative) and (2) distractor condition (3 levels: NoDis, DIS1 and DIS2) on median reaction times (RTs) of correct responses and on percentage of incorrect responses was tested using a linear mixed-effects models using the lme4 package (Bates et al. 2014) for R (Team 2014). For post-hoc analysis we used the Lsmean package (Searle et al. 1980) where p-values were considered as significant at p<0.05 and adjusted for the number of comparisons performed (Tukey method).

### 1.4 Brain Signal Recordings

Simultaneous EEG and MEG data were recorded, although the EEG data will not be presented here. The MEG data were acquired with a 275-sensor axial gradiometer system (CTF Systems Inc., Port Coquitlam, Canada) with continuous sampling at a rate of 600Hz, a 0–150Hz filter bandwidth, and first-order spatial gradient noise cancellation. Moreover, eye-related movements were measured using vertical and horizontal EOG electrodes. Head position relative to the gradiometer array was acquired continuously using coils positioned at three fiducial points; nasion, left and right pre-auricular points. Head position was checked at the beginning of each block to control head movements.

In addition to the MEG/EEG recordings, T1-weighted three-dimensional anatomical images were acquired for each participant using a 3T Siemens Magnetom whole-body scanner (Erlangen, Germany). These images were used for reconstruction of individual head shapes to create forward models for the source reconstruction procedures. The processing of these images was carried out using CTF’s software (CTF Systems Inc., Port Coquitlam, Canada).

### 1.5 Data Pre-processing

Only correct trials were considered for electrophysiological analyses. Data segments for which the head position differed for more than 10 mm from the median position during the 10 blocks were excluded. In addition, data segments contaminated with muscular activity or sensor jumps were excluded semi-manually using a threshold of 2200 and 10000 femtoTesla respectively. For all participants, more than 75 % of trials remained after rejection for further analyses.

Independent component analysis was applied on the band-pass filtered (0.1-40Hz) data in order to remove eye-related (blink and saccades) and heart-related artefacts. Subsequently, components (four on average) were removed from the non-filtered data via the inverse ICA transformation. Data were further notch filtered at 50, 100 and 150Hz and high-pass filtered at 0.2Hz.

### 1.6 Distractor-related Gamma Activity

#### 1.6.1 Gamma Band (Sensor level): Definition

The goal of this step was to define the time-frequency range of gamma activity of interest. First, for each distractor onset time-range, surrogate distractors were created in the NoDIS trials with similar distribution over time than the real distractors. Afterwards, the time-frequency power, of distractor (and of surrogate distractor) trials was calculated using Morlet Wavelet decomposition with a width of four cycles per wavelet (m=7; Tallon-Baudry and Bertrand 1999) at center frequencies between 40 and 150 Hz, in steps of 1 Hz and 10 ms. Activity between 0 and 0.35s post-distractor onset and 50-110Hz was contrasted between distractor and surrogate trials using a nonparametric cluster-based permutation analysis (Maris and Oostenveld 2007). This contrast extracts distractor-related activity clear of cue-related activity (see Figure 2).

**Figure 2.**
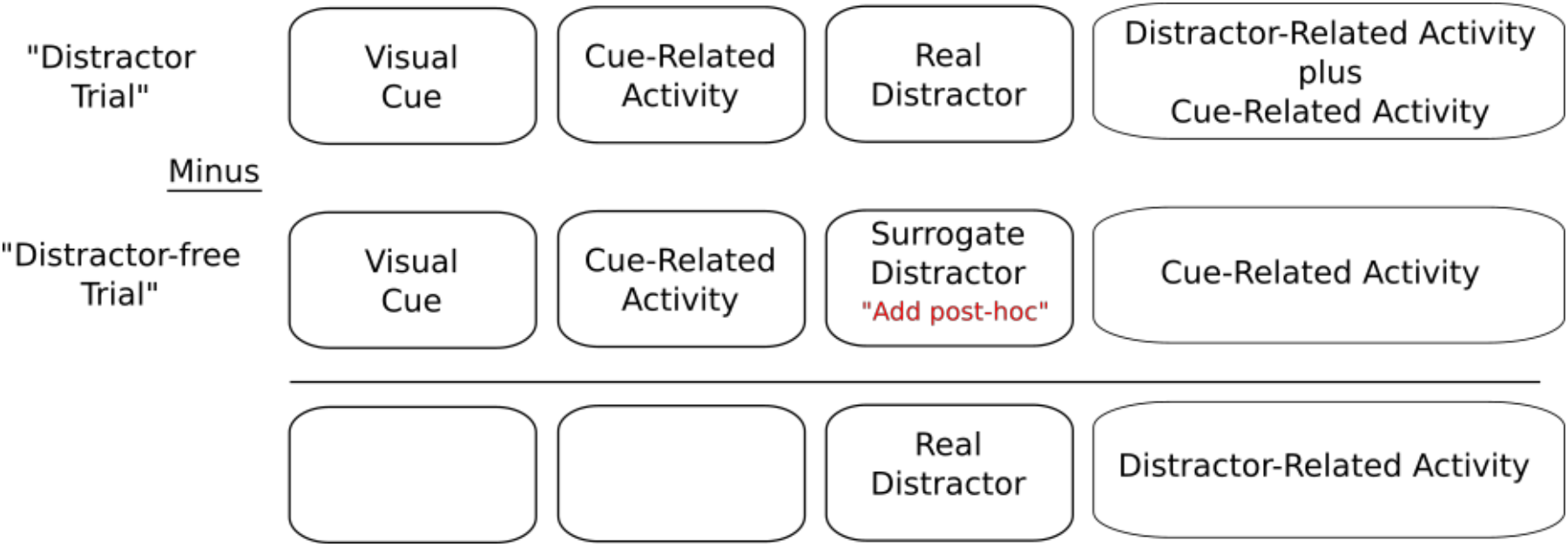
Schematic depiction for baseline correction of trials with a distracting sound.

#### 1.6.2 Gamma Band (Source level): Computation

The goal of this step was to estimate the brain regions driving gamma activity in response to distracting sounds in the time-frequency window of interest (0.1-0.3 s post-distractor onset, 60 to 100Hz) defined from the sensor-level analysis (part 1.6.1). We utilized the frequency– domain adaptive spatial technique of dynamical imaging of coherent sources (DICS; Gross et al., 2001). Data from both surrogate and real distractors were concatenated, and cross-spectral density (CSD) matrix (−0.2 to 0.6 s, relative to real/surrogate distractor onset) were calculated using the multitaper method with a target frequency of 80 (±30) Hz. For each participant, an anatomically realistic single-shell headmodel based on the cortical surface was generated from individual head shapes (Nolte, 2003). A grid with 0.5 cm resolution was normalized on an MNI template, and then morphed into the brain volume of each participant. Leadfields for all grid points along with the CSD matrix were used to compute a common spatial filter that was used to estimate the spatial distribution of power for time-frequency window of interest (0.1-0.3 s post-distractor onset, 60 to 100Hz).

#### 1.6.3 Gamma Band (Source Level): Analysis

For each participant, we estimated source-level activity (0.1-0.3 s post-distractor onset, 60 to 100Hz) for each cue category (informative and uninformative) and for both cue categories concatenated. We performed two analyses:

1. In order to characterize the brain areas activated in the gamma band during the distracting sound presentation, distractor-related gamma activity was contrasted to surrogate distractor-related gamma activity
2. In order to investigate the interaction between distractor response and cue information, surrogate-corrected gamma activity in the informative cue condition was contrasted to surrogate-corrected gamma activity in the uninformative cue condition.

Both tests have been carried out using non-parametric cluster-based permutation analysis (Maris and Oostenveld 2007). Please note, that for these tests, cluster permutations control for multiple comparisons in the source space dimension.

#### 1.6.4 Gamma Band (Source Level): Connectivity Analysis

The aim of this analysis was to identify the brain regions that could be functionally connected in the gamma band to the auditory cortices during the presentation of the distracting sound. We have extracted the complex values containing phase information into source space using partial canonical coherence (PCC) beamformer, a computationally efficient alternative to the DICS that provides the possibility of extracting both power and phase information on the source level. For each participant:

1. Similarly, to the DICS beamformer, data from both surrogate and real distractors were concatenated, and cross-spectral density (CSD) matrix (−0.2 to 0.6 s, relative to real/surrogate distractor onset) were calculated using the multitaper method with a target frequency of 80 (±30) Hz. Leadfields for all grid points along with the CSD matrix were used to compute a common spatial filter that was used to estimate the spatial distribution of power and phase for the time-frequency window of interest (0.1-0.3s post-distractor onset, 60 to 100Hz).
2. One auditory region of interest (ROI) was defined as including, in both hemispheres, the Broadmann areas 22, 41 and 42, according to the Talairach Tournoux atlas (Talairach and Tournoux 1988; Lancaster et al. 1997).
3. Phase synchrony (Lachaux et al. 1999) between each voxel in the auditory ROIs and all other voxels was calculated, averaged across voxels of the auditory ROI, and then Fisher Z transformed. Thus, for each extra-auditory brain voxel, a single phase synchrony value with the entirety of the auditory ROI was obtained.

Finally, distractor-related gamma phase synchrony was contrasted to surrogate distractor-related gamma phase synchrony using non-parametric cluster-based permutation analysis (Maris and Oostenveld 2007). Please note, that for this test, cluster permutations control for multiple comparisons in the source space dimension.

## 2 Results

### 2.1 Behavioral Analysis

Participants correctly performed the CAT in 96.04 ± 0.29 SEM % of the trials. The remaining trials were either incorrect trials (3.95 ± 0.29 SEM %), missed trials (0.27 ± 0.06 %) or trials with FAs (0.02 ± 0.01 %).

#### 2.1.1.1 Median Reaction Times

We found a significant main effect of cue category (F(1, 20) = 4.9, p = 0.02, η^2^ = 0.36) on median reaction times in correct trials. Participants were faster when the cue was informative in comparison to the uninformative cue. In addition, we found a significant main effect of the distractor condition (F(2, 42) = 34.2, p < 0.01, η^2^ = 0.49). Post-hoc tests indicated that, in comparison to the NoDIS condition, participants were faster in the early DIS1 condition (p < 0.001) but slower in the late DIS2 condition (p < 0.001). Participants were also faster in the early DIS1 than in the late DIS2 condition (p < 0.001). No interaction effect was found significant.

#### 2.1.1.2 Percentage of Incorrect Responses

Only a main effect of the distractor condition (F(2, 42) = 3.8, p = 0.02, η^2^ = 0.17) was found significant on the percentage of incorrect responses. Post-hoc tests indicated that participants, committed more errors in the late DIS2 condition in comparison to the NoDIS (p = 0.02) and early DIS1 (p = 0.07) conditions.

**Figure 3.**
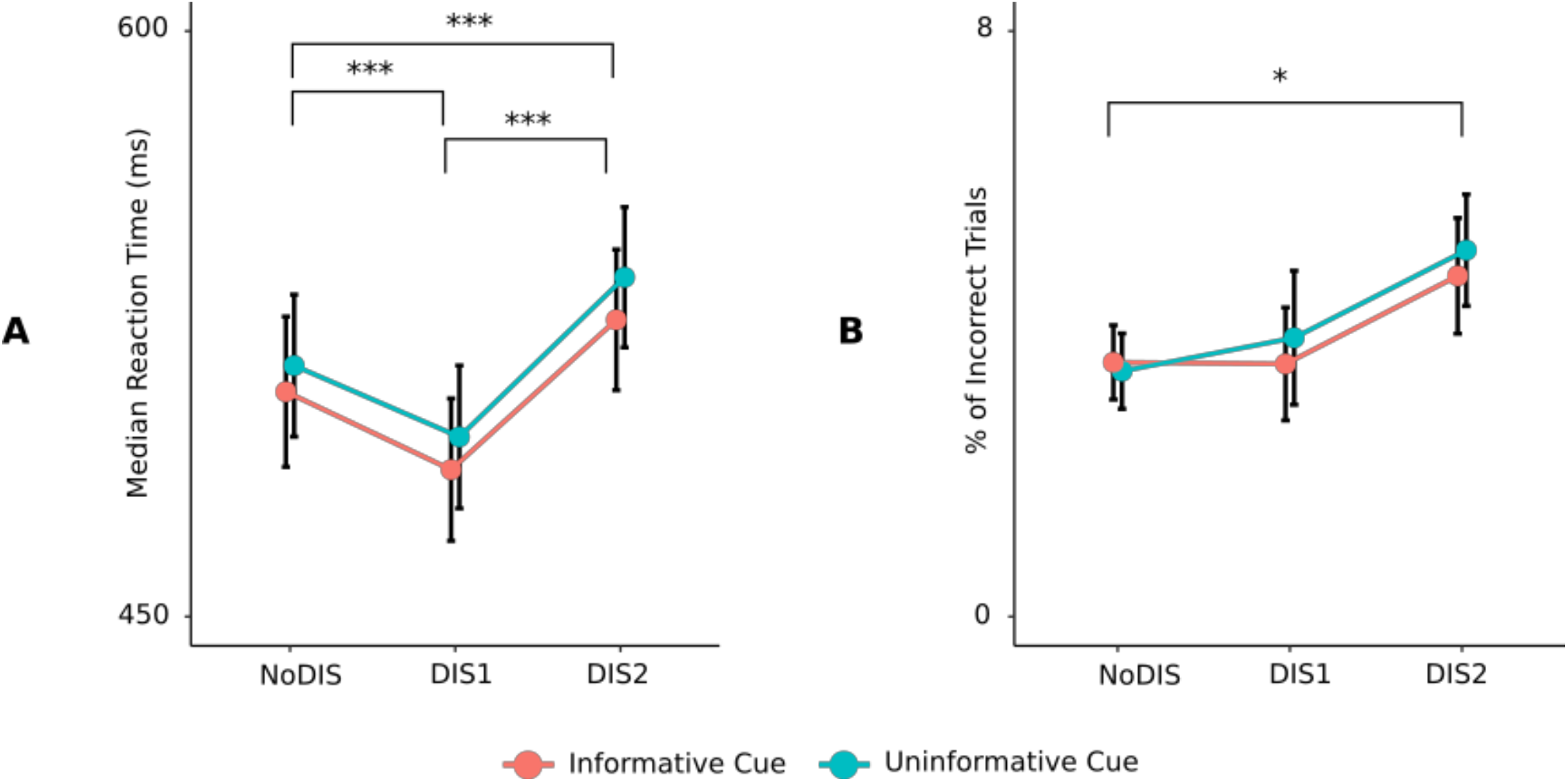
**A.** Median Reaction Times (RTs) according to cue and distractor conditions. **B.** Percentage of Incorrect Responses according to cue and distractor conditions. * *P* < 0.05, ** *P* < 0.01, *** *P* < 0.001. Error bars represent SEM.

### 2.2 Gamma Activity Analysis

#### 2.2.1 Gamma Sensor Level Activation

Real-distractor high-frequency activity was contrasted to that of surrogate-distractor using non-parametric cluster-based permutation testing. As shown in Figure 4, this contrast revealed one significant positive cluster (p = 0.002) indicating an increase in gamma activity centered spatially around left and right temporal sensors (Fig 4A), temporally between 0.1 and 0.3s post-distractor onset (Fig 4B & C), and frequency-wise, between 60 and 100 Hz (Fig 4B & D).

**Figure 4.**
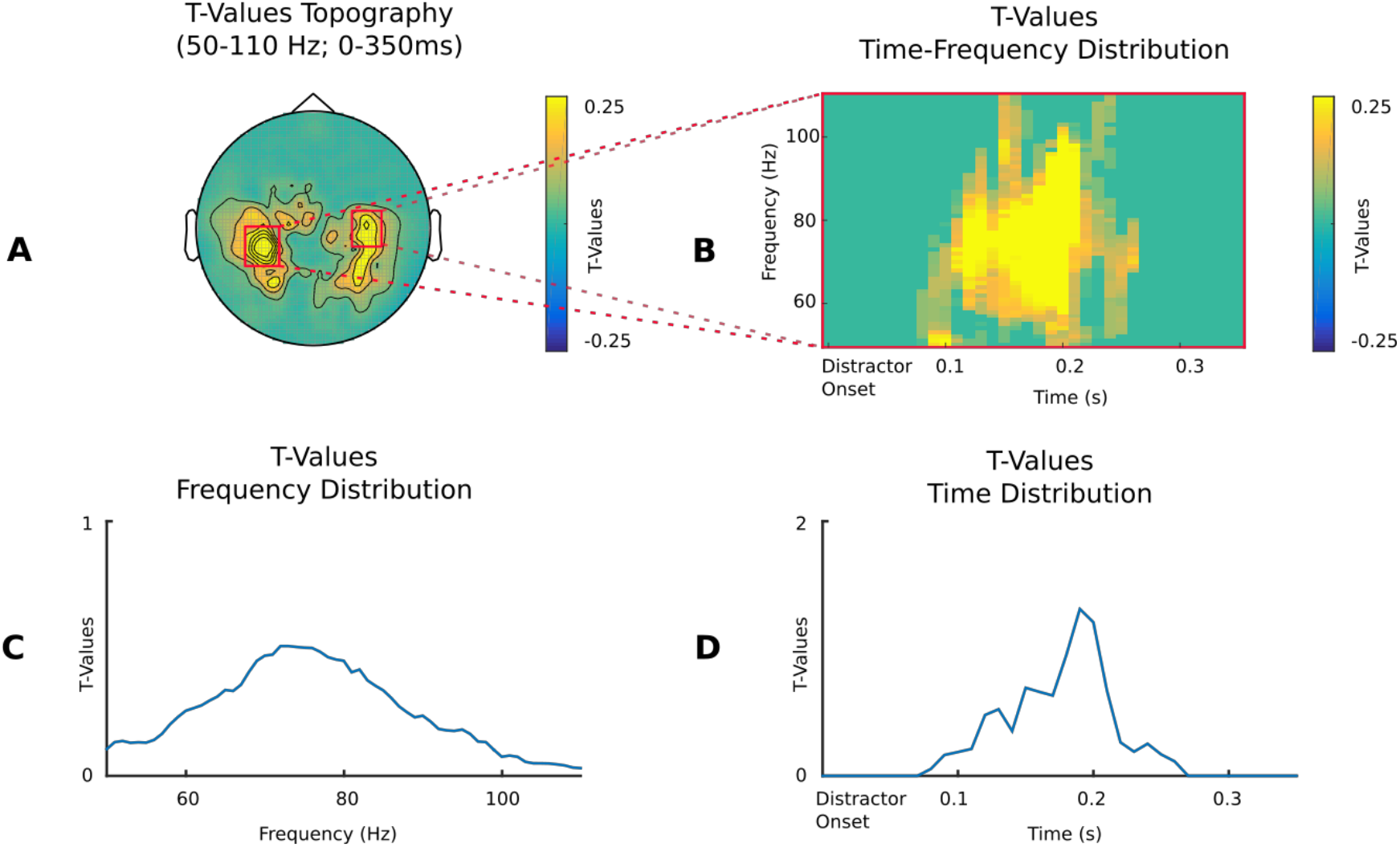
Gamma activity to distractor at the sensor level. **A.** Topographical maps averaged between 50-110 Hz and 0-0.35 seconds post-distractor onset, of the t-values, masked at p < 0.05 of the contrast between distractor and surrogate distractor gamma activity. **B.** Time-frequency representations of the t-values (of the aforementioned test) of the sensors highlighted by red boxes in A. **C.** Frequency distribution of t-values of the aforementioned sensors averaged across the time dimension. **D.** Time distribution of t-values of the aforementioned sensors averaged across the frequency dimension.

#### 2.2.2 Gamma Source Level Activation

The aim of this test was to highlight the regions driving gamma activity observed at the sensor level. Based upon the sensor level results, we have computed DICS beamformer sources for each participant between 60-100Hz and 0.1-0.3s post-distractor. Real-distractor gamma activity was contrasted to that of surrogate distractors using non-parametric cluster-based permutation testing. A significant positive cluster (p < 0.01) indicating an increase in gamma activity notably in (1) the left and right auditory cortices, (2) the left and right tempo-parietal junctions, and (3) the right ventrolateral prefrontal cortex (vlPFC). Other regions included the calcarine, the anterior, middle and posterior cingulate gyri, the inferior temporal gyri and the Precuneus.

**Figure 5.**
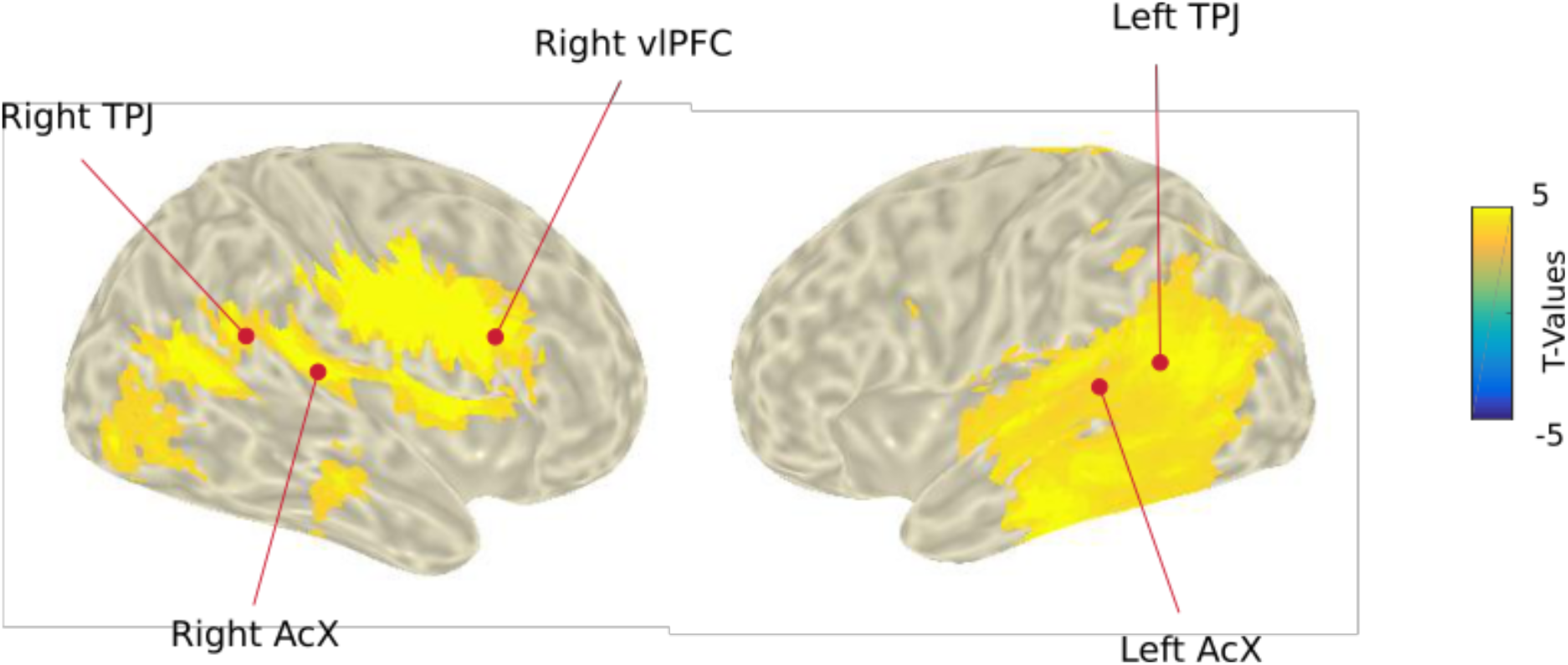
Gamma activity to distractor at the source level. Distributions of T-values, masked at *p<0.05*, from Cluster Based Permutation tests contrasting real and surrogate distractor gamma activity (60-100Hz and 0.1-0.3 post-distractor) at the source level. AcX: Auditory Cortex. TPJ: temporo-parietal junction. vlPFC: ventrolateral prefrontal cortex.

#### 2.2.3 Gamma Source Level Cue Effect Comparison

To investigate the effect of top-down attention on bottom-up processing, we analysed the effect of cue information on the gamma response to distracting sounds. For each participant, real distractor source-level data (60-100 Hz, 0.1-0.3s) were baseline corrected by subtracting surrogate-distractor activity. Corrected distractor gamma activity was contrasted between the two cue categories (informative vs. uninformative) using non-parametric cluster-based permutation testing. A significant cluster (p = 0.014) extended notably to (1) the left dorsomedial and ventromedial prefrontal cortices, and (2) the left anterior cingulate gyrus. Other regions included the left pre- and post- central gyri, the left supplementary motor area, the left superior parietal lobule. All these regions displayed a significantly higher activation when the distracting sound was preceded by an informative cue rather than an uninformative cue.

**Figure 6.**
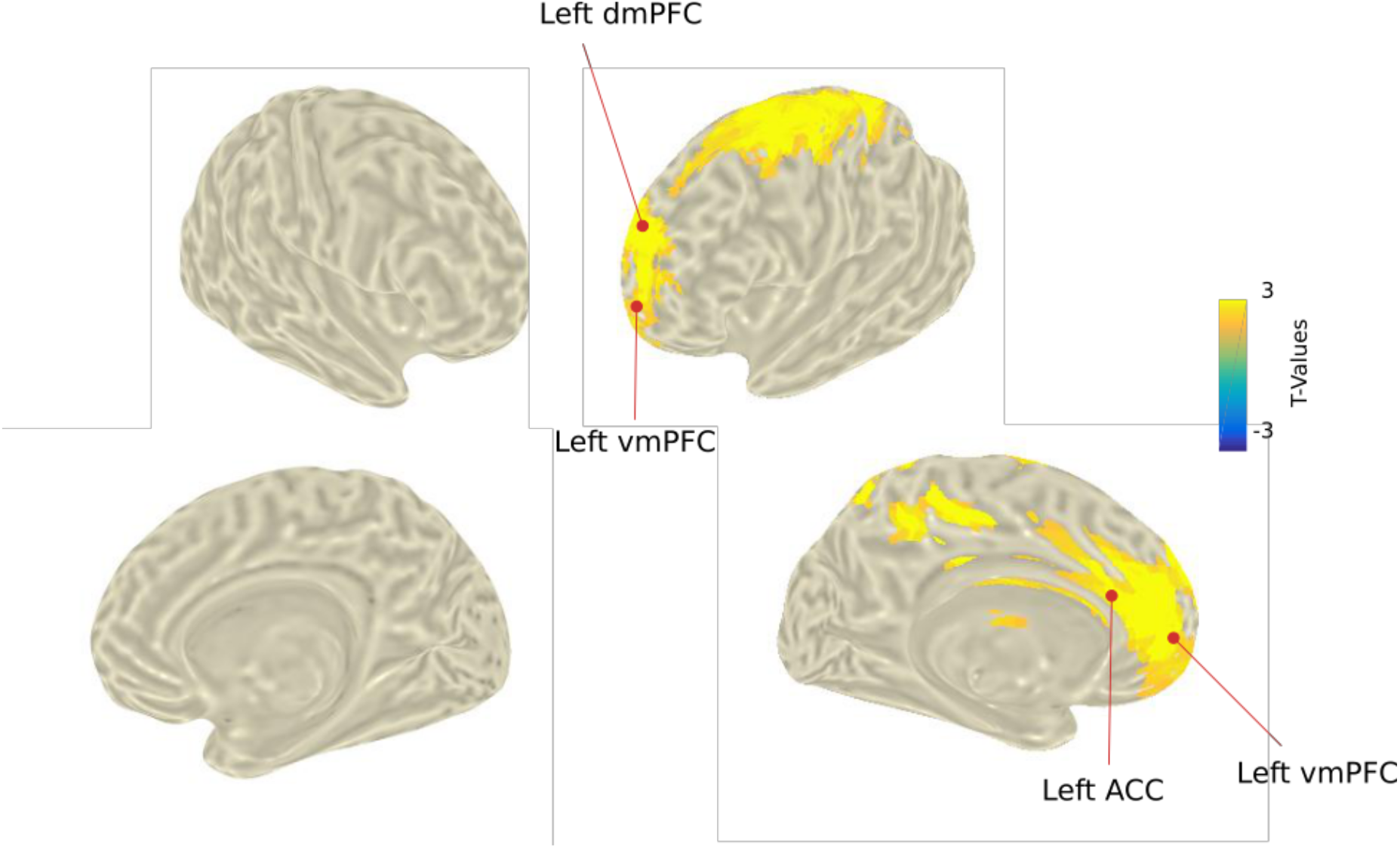
Top-down modulation of gamma activity to distractor at the source level. Distributions of T-values, masked at *p<0.05*, from Cluster Based Permutation tests contrasting surrogate-corrected distractor gamma activity (60-100Hz and 0.1-0.3 post-distractor) between informative and uninformative cue conditions. dmPFC: dorsomedial prefrontal cortex. vmPFC: ventromedial prefrontal cortex. ACC: anterior cingulate cortex.

#### 2.2.4 Gamma Source Level Connectivity Analysis

Real-distractor phase synchrony (both auditory ROIs averaged being the reference) in the gamma band (60-100 Hz, 0.1-0.3s) was contrasted to that of surrogate distractors using non-parametric cluster-based permutation testing. A significant cluster (p < 0.01) indicated an increase in gamma synchrony (during the presentation of distracting sounds) between the auditory cortices and the ventrolateral and dorsolateral prefrontal cortex in both hemispheres. Other regions included the pre- and post- central gyri, the supplementary motor areas, the frontal eye fields and the paracentral lobule, in the left hemisphere.

**Figure 7.**
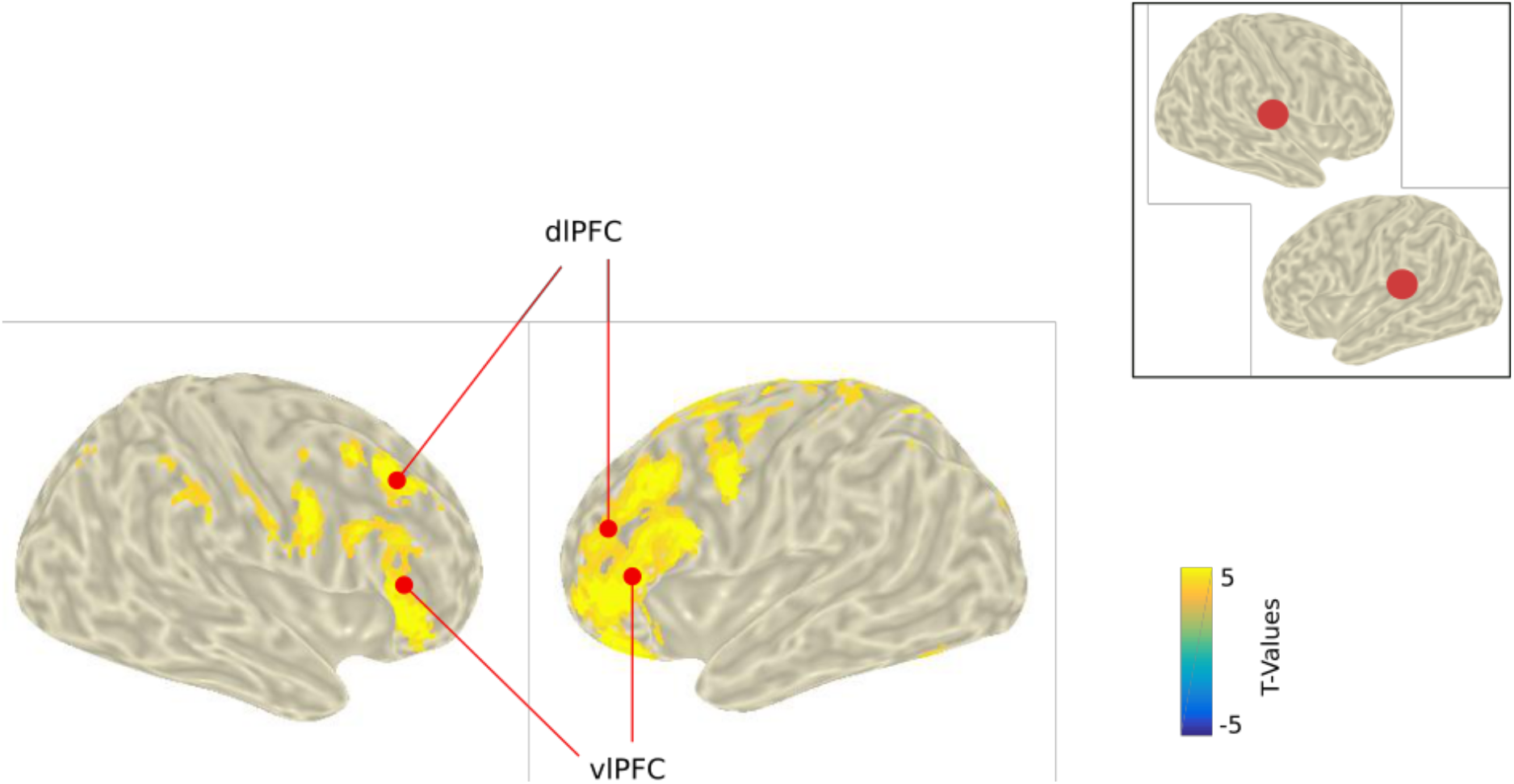
Gamma connectivity to distractor at the source level. Distributions of T-values, masked at *p<0.05*, from Cluster Based Permutation tests contrasting real and surrogate distractor gamma phase synchrony (60-100Hz and 0.1-0.3 post-distractor) at the source level between the auditory ROI (bilateral auditory cortices, see image in the right top corner) and all other cortical voxels. dlPFC: dorsolateral prefrontal cortex. vlFPC: ventrolateral prefrontal cortex.

## 3 DISCUSSION

In the present study, we have demonstrated that in response to salient unexpected distracting sounds, the auditory cortices and the temporo-parietal junctions in both hemispheres, and the right ventrolateral PFC were activated in the gamma band. In addition, modulation by top-down attention of gamma activity to distracting sounds was found in the left dorsomedial and ventromedial prefrontal cortices. Finally, we have evidenced synchrony in the gamma band between regions in the dorsolateral and ventrolateral prefrontal cortices and the auditory cortices during distracting sound processing.

### 3.1 Behavioral measures of TD and BU attentional mechanisms

Behaviorally, participants discriminated the target pitch faster in trials with an informative cue in comparison to trials with an uninformative cue. This effect is in agreement with several previous studies (Posner 1978; ElShafei et al. 2018). It is most likely related to differences in TD anticipatory attention since the informative cue provided additional information solely about the location of the target and not about its category neither its mapped response, leading to equivalent motor preparation across conditions.

In trials with distracting sounds, participants responded faster to the following target in trials with early distracting sounds rather than with late distracting sounds. This pattern can be explained in light of the phenomena triggered by a distracting sound (review in Näätänen 1992; Bidet-Caulet et al. 2014; Masson and Bidet-Caulet 2018): (1) a persistent increase in arousal resulting in RT reduction (behavioral benefit) and (2) a strong transient attentional capture (exogenous orienting) leading to RT augmentation (behavioral cost); with the behavioral net effect of distracting sound varying according to the time interval between the distracting and the target sounds. Importantly, we also found that participants were less accurate to discriminate the auditory targets when preceded by late distracting sounds in comparison to the no distractor condition. This result provides further evidence of a detrimental behavioral effect of distracting sounds which orient attention away from the task at hand.

Therefore, these behavioral results demonstrate that, in the CAT paradigm, TD attention is enhanced in trials with informative cue, and that a strong transient bottom-up attentional capture is triggered by distracting sounds.

### 3.2 Activation of the Ventral Bottom-Up Attentional Network in the Gamma band

In line with our hypothesis, in response to an unexpected salient distracting sound, we observed an increase in gamma activity in the left and right auditory cortices, the bilateral temporo-parietal junctions, and the right ventrolateral prefrontal cortex. This present result is highly consistent with the proposal by Corbetta and Shulman (2002; 2008), based on functional magnetic resonance imaging (fMRI) studies of visual attention, that the ventral attention system, involved in bottom-up attention, includes the temporo-parietal junction and the ventral prefrontal cortex. This present finding is also in agreement with fMRI (e.g. Salmi et al. 2009; Alho et al. 2014; Salo et al. 2017) and MEG/EEG (e.g. Ahveninen et al. 2013) studies in the auditory modality. In addition, the right lateralization of the frontal activation is consistent with fMRI findings of a ventral network predominantly localized to the right hemisphere in the visual modality (Corbetta and Shulman 2002; Corbetta et al. 2008). Also, the bilateral activation of the TPJ is in line with a stronger activation of the left TPJ to auditory than to visual irrelevant oddball stimuli as shown in a meta-analysis of fMRI studies (Kim 2014).

Importantly, we found that the gamma activation in the ventral network in response to distracting sounds lasted from 100 to 300ms after distracting sound onset. This result confirms that BU attentional capture is a rapid and transient phenomenon, in agreement with behavioral costs observed only for late distracting sounds offsetting between 50 and 350 ms before target onset. Such activation in high frequency gamma oscillations is also consistent with the fast nature of the attentional capture phenomenon. Finally, it is worth noting that the activation of the ventral attentional network was found in response to entirely task-irrelevant distracting sounds, contrary to earlier studies which suggested that task-relevance rather than saliency is critical to the engagement of the ventral network (Serences et al. 2005; Indovina and Macaluso 2007; Corbetta et al. 2008). The short duration of the ventral network activation might have precluded its observation using techniques with low temporal resolution (fMRI) as previously proposed by Chica and colleagues (2013).

### 3.3 The PFC and the balance between TD and BU attentional mechanisms

We did not find any significant effect of cue information on gamma activity within the regions of the ventral attentional network in the first 300 ms. However, in several regions of the left prefrontal cortex, notably (and contrary to our original hypothesis) in the dorso- and ventro-medial prefrontal cortices and the anterior cingulate cortex (ACC), gamma activity was more pronounced in response to distracting sounds preceded by an informative cue rather than an uninformative cue.

These medial prefrontal regions have been hypothesized to play a role in the inhibition of irrelevant stimuli. Indeed, in humans, an increased BOLD activation of these regions is observed during presentation of irrelevant salient stimuli (Salmi et al. 2009); and an enhanced P3 response to such stimuli is found after damage of the medial PFC (Rule et al. 2002). In the non-human primate auditory system, such role is supported by cortico-cortical connections between these regions (the medial PFC and the ACC) and inhibitory neurons in auditory association regions in order to suppress irrelevant signals (Matsumoto and Tanaka 2004; Barbas et al. 2005, 2012; Medalla et al. 2007). Importantly, electrical stimulation of the ACC has been shown to reduce auditory evoked activity in non-human primate superior temporal cortices (Müller-Preuss et al. 1980; Müller-Preuss and Ploog 1981), providing direct evidence of an inhibitory role of these medial prefrontal regions. Therefore, in the present study, the larger gamma activation of the medial PFC regions and the ACC, during distracting sounds preceded by an informative cue, could reflect a strong and fast inhibitory signal to regions involved in the processing of task-irrelevant information. This stronger inhibition could result from an increased top-down attention load with informative cue, in line with shorter reaction times, compared to trials with uninformative cue.

The medial prefrontal localization of such regions contradicts our original hypothesis that more lateral prefrontal regions would play a prominent role in orchestrating the interplay between TD and BU attentional mechanisms, as suggested by previous studies using fMRI in human subjects (Fox et al. 2006; Corbetta et al. 2008; Alho et al. 2014; Katsuki and Constantinidis 2014; Vossel et al. 2014). However, we believe that such role for the lateral PFC cannot be ruled out. As evidenced in the non-human primate brain, the ACC has excitatory and inhibitory connections to the anterior and posterior parts of the lateral PFC, respectively (Medalla and Barbas 2010), with the former momentarily suspending current tasks and the latter facilitating attentional switch to a novel task (review in Barbas et al. 2012). In the context of the present paradigm, a stronger ACC to lateral PFC signal could facilitate switching from the TD task to the BU distracting sound processing, and vice versa. The opposite effects on the anterior and posterior parts of the lateral PFC combined to the insufficient spatial resolution of MEG could preclude the observation of a significant increase in gamma activation in the lateral PFC. Therefore, the medial PFC could support the interplay between top-down and bottom-up attention by (1) exercising a TD inhibitory attentional control via direct projection to the auditory cortices, and (2) by controlling task-switching between TD and BU brain operations via projections to the lateral PFC. Importantly, in the present study an increase in gamma synchrony was found, during the presentation of a distracting sound, between the auditory cortices and several distant brain regions, notably the dorso- and ventro-lateral prefrontal cortices, but not the medial prefrontal regions. This result argues for a strong implication of the lateral PFC.

Activation of the lateral PFC has also been associated with distractor suppression (Dolcos et al. 2007; Suzuki and Gottlieb 2013). It has been demonstrated that reversible inactivation of the lateral PFC (specifically the dorsolateral prefrontal cortex) results in increased distractibility in monkeys (Suzuki and Gottlieb 2013), and its stimulation with tDCS decreases susceptibility to attentional capture in humans (Cosman et al. 2015). Thus, the lateral PFC could be attributed a role in cognitive inhibitory control. We posit that the lateral PFC, through gamma phase synchrony, could play a role in the propagation of the top-down inhibitory signal from the medial PFC and ACC to the auditory cortices, in order to filter out task-irrelevant distracting sounds in anticipation of the upcoming relevant target sound.

This interpretation is in line with the Communication-Through-Coherence (CTC) hypothesis which proposes that anatomical connections are dynamically rendered effective or ineffective through the presence or absence of rhythmic synchronization, particularly in the gamma band (Fries 2005). Moreover, this suggested link between the lateral and medial subdivisions of the prefrontal cortex is in line with previous studies highlighting high interconnectivity between the lateral and medial PFC (Miller and Cohen 2001; Cole et al. 2013). In their study, Cole and colleagues (2013) demonstrated that the lateral PFC along with the posterior parietal cortex, constitute a highly flexible connectivity hub that could be involved in implementing task demands by biasing information flow across multiple large-scale functional networks. Thus, the lateral PFC could act as an inhibitory (control) signal relay hub between nodes of the ventral bottom-up attentional network (e.g. the auditory cortex) and the medial PFC and ACC. Consequently, the lateral PFC could orchestrate the interplay between dorsal and ventral attentional networks, by relaying inhibitory signal from the medial PFC, leading to switching between TD attention towards the upcoming target and BU attention to the distracting sound.

### 3.4 Conclusion

Using the high temporal and spatial resolution of magnetoencephalography, we demonstrate for the first time, in the human brain, how gamma activity would support activation and communication within the ventral BU attentional network, as well as the interaction between BU and TD attention. This corroborates the proposed role for gamma oscillations as a promoter of rapid transfer of information through the cortical hierarchy (Sedley and Cunningham 2013). Moreover, this finding fits in a wider framework proposing that activity in different attentional networks would be supported by different frequency bands. More precisely, slow oscillations (namely alpha) would support more top-down attentional mechanisms, while faster activities (namely gamma) would support more bottom-up attentional mechanisms (Buschman and Miller 2007). Importantly, the present results strongly suggest that the medial prefrontal cortex would control the balance between top-down and bottom-up mechanisms of attention, while the lateral prefrontal cortex would orchestrate the interaction between top-down, bottom-up, and inhibition attentional networks.

## 4 ACKNOWLODGEMENTS

We would like to thank Sebastien DALIGAULT and Claude DELPUECH for technical assistance with the acquisition of electrophysiological data. This work was supported by the French National Research Agency ANR-14-CE30-0001-01 (ABC), by the Rhone-Alpes Region (HE) and by the LABEX CORTEX (ANR-11-LABX-0042). This work was performed within the framework of the LABEX CELYA (ANR-10-LABX-0060) of Université de Lyon, within the program ‘‘Investissements d’Avenir’’ (ANR-11-IDEX-0007) operated by the French ANR.

